# Bacterial community composition of recycled irrigation water of a NFT-experimental system, with and without a slow sand filter

**DOI:** 10.1101/486464

**Authors:** Giovanni Cafà, Richard Thwaites, Matthew J Dickinson

## Abstract

The bacterial community composition (BCC) of recycled irrigation freshwater was monitored on a Nutrient Film Technique (NFT)-type experimental hydroponic system. Two identical NFT systems in a greenhouse were used to grow tomato plants. One was connected to a slow sand filter (SSF), and one was used as control. DNA was isolated from irrigation freshwater, and molecular methods were carried out to characterise the BCC. These included terminal-restriction fragment length polymorphism (T-RFLP), cloning and 454 pyrosequencing. A total number of 291 442 trimmed sequences, of 211 bp average length. A strong differentiation of the bacterial community composition in recycled irrigation water was triggered by the activity of the SSF. Phylogenetic affiliation revealed that *Bacilli, Alpha-* and *Gammaproteobacteria*, and *Nitrospira* were differentially abundant during filtration with the SSF. This study showed that SSF modified the relative amount of a set of bacterial genera. These included *Pseudomonas, Bacillus, Flavobacterium, Burkholderia* and *Azospirillum*. These bacteria have been previously described as plant growth promoting rhizobacteria (PGPR). The presence, increased or exclusive, in Water *ssf* of bacteria such as *Bacteroidetes, Gemmatimonadetes, Nitrospira, Firmicutes, Alpha-, Beta-, Gamma-* and *Deltaproteobacteria*, was associated with an increased biomass in plants. Such findings are promising for future applications of a combined system NFT-SSF: NFT guarantees a controlled closed environment for the growth of plants, while the SSF secures the microbiological balance of recycled irrigation water.

## INTRODUCTION

Recycled irrigation freshwater is an emerging agroecosystem of growing interest. In recent years, the possibility of growing plants using re-circulated and replenished nutrient solution has been increasingly investigated worldwide (Garland, 1994; Zhang and Tu, 2000; Alsanius *et al*., 2001; Postma *et al*., 2001; Frenkel *et al*., 2010). It has been demonstrated that the same yields can be obtained in closed systems (where surplus solution is recovered, replenished, and recycled) as compared to open systems (once the nutrient solution is delivered to the plant roots, it is not reused) (Jensen, 1997), as long as good hygienic and environmental conditions are maintained (Gertsson *et al*., 1994). The use of recycled irrigation freshwater can effectively reduce water usage and mitigate nutrient runoff from nursery production sites. However, serious concerns exist regarding the spread of phytopathogenic microorganisms via the recycled water (Waechter-Kristensen *et al*., 1997; Pagliaccia *et al*., 2008).

Irrigation freshwater can be recycled with nutrient film technique (NFT). NFT is a simple and cost effective method for growing plants, where roots are emerged in a continuous flow of re-circulating water containing all the nutrients that plants need. A root mat develops partly in the shallow stream of re-circulating water and partly above it. Thus the stream is very shallow, and the upper surface of the root mat that develops is above the water (Cooper, 1979). NFT can save water and reduce pollution associated with the need to discharge used solutions into the environment (Calvo-Bado *et al*., 2006).

Recycling the water supply used for irrigation can generate contamination with plant pathogens from several sources. The pathogen may be a natural inhabitant of the water source, or reside in the soil, or in infected resident plants near the water and only be a transient inhabitant of the irrigation water (Hong and Moorman, 2005). Any infectious propagule has the potential to make contact with the irrigation water and, upon entry into the nutrient solution, can ultimately make contact with a root (Stanghellini and Rasmussen, 1994). This contact, which is possible at several points along the distribution path (Hong and Moorman, 2005), can create a high risk of contamination for the entire crop. NFT systems are particularly exposed to the interaction between water and plant roots because of the density of roots exposed in the air (Stanghellini and Rasmussen, 1994; Clematis *et al*., 2009). The layout of the NFT method, in which the plants are produced with bare roots, puts the crop in danger and, despite the many advantages introduced by the use of recycled irrigation freshwater, can create the condition for contamination by pathogens (Hong and Moorman, 2005).

Fungi are not the only problem related to recycled irrigation freshwater. The establishment of bacterial pathogens also represents a concern when producing minimally processed ready-to-eat vegetables such as tomatoes and lettuce. *Enterobacteriaceae* such as *Escherichia coli* and *Salmonella* spp. are common in greenhouse production, and the use of soil-less technologies may increase apprehension. Viruses can also spread via irrigation water after being released by plant roots (Büttner *et al*., 1995): Pelargonium flower break virus can spread in recirculating nutrient solutions in greenhouses (Berkelmann *et al*., 1995) as can tomato mosaic virus (Pares *et al*., 1992).

Risk of contamination of irrigation freshwater can, however, be reduced by the use of disinfection methods such as slow sand filtration (SSF). SSF is a disinfection method available since 1974 (Huisman and Wood, 1974, Calvo-Bado et al., 2003). It is a technique that involves the slow passage of water or liquid through a porous medium, which for horticultural uses is commonly constituted of sand. This technology is employed to reduce contaminants from freshwater, as a result of a complex consortium of microorganisms that develops and interacts in the top layer of the sand column, called *schmutzdecke*. The schmutzdecke – a biologically active layer, can generally be considered as a gelatinous biofilm containing a consortium of bacteria, fungi, algae, protozoa, rotifers, and a range of aquatic insect larvae. The activity of the microbiota of the schmutzdecke is directly responsible for much of the treatment function, although it should be stressed that the underlying sand or other bed material is also responsible for the removal of various fractions from the water. A ripening period of 3-6 weeks is required for this layer to form, during which the filter performance is sub-optimal (Joupert and Pillay, 2008).

Bacteria are an important component of the schmutzdecke in SSF. Bacterial communities in the schmutzdecke have been studied since 1952 (Calaway *et al*., 1952), where dominant organisms were identified with traditional culture dependent techniques. One of the first studies that applied modern molecular techniques to understand the microbiology of SSF was conducted by Petri-Hansen *et al*. (2006). The bacterial population in the filter was found to be more diverse in distribution between taxonomic groups and of different physiological functions than previously recognised. A large component of the community was comprised of *Proteobacteria* (25%), while the remaining 75% was of less commonly encountered bacterial taxa. Petri-Hansen *et al*. (2006) showed that the bacterial composition of the schmutzdecke was affected by influent temperature, and mainly determined by the autochthonous bacteria from the aquatic environment. However, a more detailed understanding of the bacterial community in SSFs is still needed (Page *et al*., 2006), as well as the effect of the SSF on the microflora of recycled irrigation freshwater. The objectives of this study were therefore to investigate the bacterial community composition (BCC) of the recycled irrigation freshwater developing in a NFT-Type experimental system, focusing on the microflora of the water matrix. This study also describes the differences between the BCC of the recycled irrigation freshwater with and without slow sand filtration, focusing on the bacterial taxa that have a higher relative abundance caused by the activity of the SSF.

## MATERIAL AND METHODS

Tomato plants on rockwool were grown on NFT-Type experimental systems, with recycled nutrient water fully recycled in the process. DNA was isolated from the water tomato pfor the investigation of the differences of the bacterial community composition (BCC) developing in the recycled irrigation freshwater. Each experiment was carried out in a 28 days cycle with 4 time points (7,14, 21, and 28 days). All data are presented as the average of three replicates of the 28 days experiments (NFT1, NFT2, and NFT3), with the exception of the pyrosequencing analysis, which was performed on one of the three replicates (NFT1).

### Nutrient Film Technique (NFT) system

Two identical replicates of a NFT-Type system were established in a glasshouse (Sutton Bonington Campus, University of Nottingham, UK). One was connected to a slow sand filter (SSF, Fig. 1), and the other one was used as the control. Fifty ml samples of recycled irrigation freshwater were collected at 4 time points (7,14, 21, and 28 days) from the two water tanks of the NFT systems, in which tomato plants were grown (*Solanum lycopersicum* cv. Alicante).

**Figure 1.**
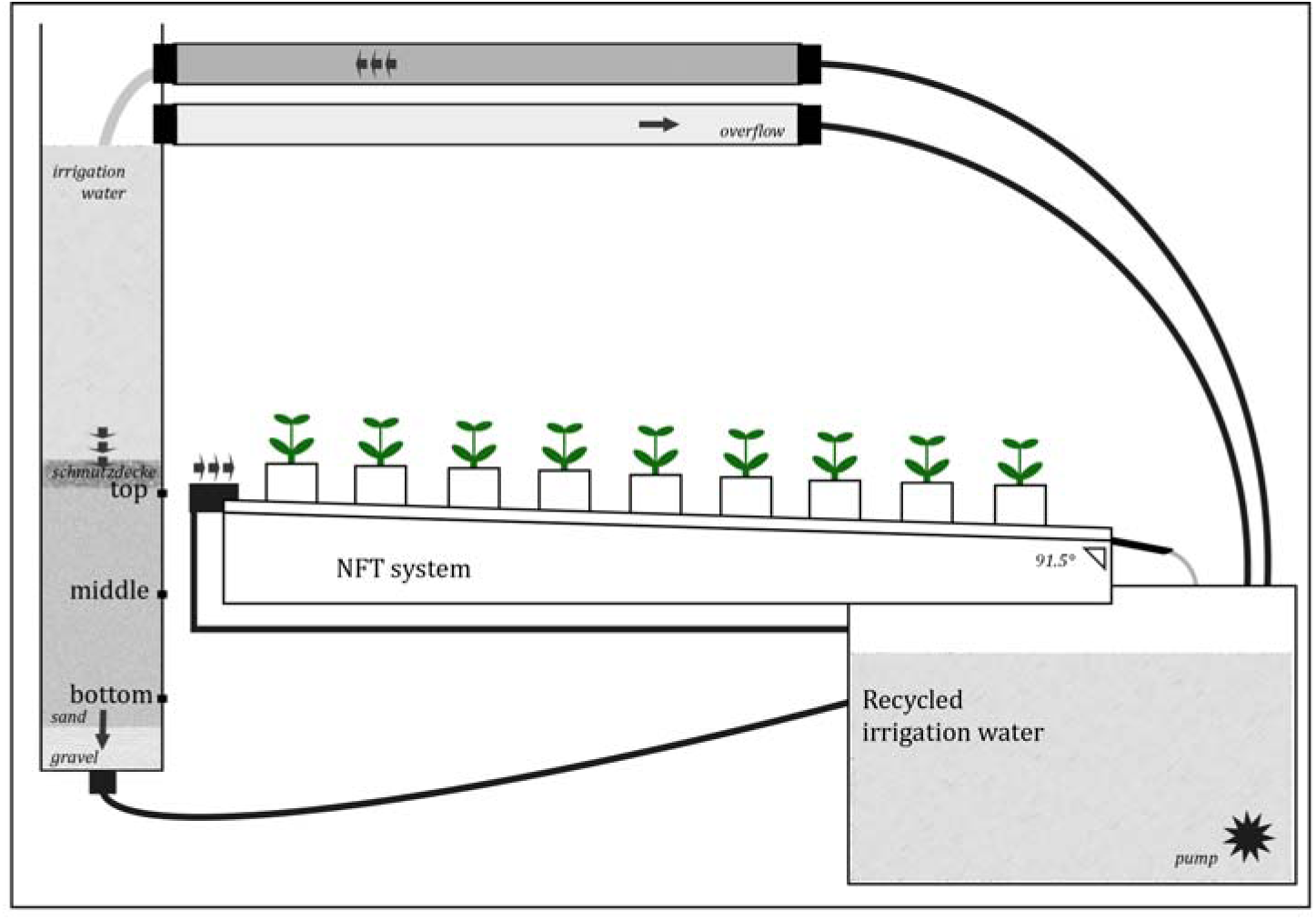
Schematic representation of the experimental NFT system connected to the slow sand filter. Recycled irrigation water was directed to the top of the column of the sand filter and to the channels of the NFT with the aid of a pump (*). Channels of PVC had an inclination of 1.5° for allowing the water to flow down the channels and returning to the tank and restart the cycle. Arrows show the direction of recycled irrigation water. The figure is not to scale.

Irrigation water was obtained from nutrient stock solution (1:200 VITAFEED-214, VITAX, Leicester, UK) added to tap water. In all the experiments, an initial volume of 100 L of irrigation water was placed in the main tank; three rows of polyvinyl chloride channels (PVC) (Geberit, Aylesford, UK) containing 9 plants each were connected to the 100 L tank, and the pump was activated to allow the continuous flow of water from the top to the bottom of the PVC channels. The PVC channels were kept at an inclination of 1.5° to allow the water to return under gravity before being recirculated by the pump.

Plants were grown from seeds of *Solanum lycopersicum* cv. Alicante (Wilkinson, Worksop, UK) in incubators for 14 days before being transferred to rockwool cubes and relocated to the NFT system. The NFT system allowed roots to develop outside rockwool cubes, and establish a thick net of roots in the PVC channel, with water continuously surrounding the roots of the plants.

The flow of irrigation freshwater through the channels was regulated at 2 L/min, and maintained at a constant rate throughout the experiments. After completion of each replicate of the experiment, the NFT-Type system was thoroughly washed before placing new plants and fresh irrigation freshwater. Each experimental replicate was carried out for 28 days. Samples of 50 ml of recycled irrigation freshwater were collected every 7 days from the tank of the control system (Water *co*) and the tank of the NFT connected to the SSF (Water *ssf*). An equal volume of irrigation freshwater was added at the time of sampling to the two tanks. This volume was variable, and depended on the consumption of water of the system, e.g. a higher volume was added in warmer months of the season.

### Characteristics of the Slow Sand Filter (SSF) connected to the NFT system

A SSF was linked to one of the two experimental NFT-Type systems (Fig. 1). The filter was prepared with a 2 m Terrain PVC pipe of 20 cm diameter mounted vertically. The bottom of the filter was filled with a 30 cm depth of gravel. The column consisted of a sand bed of 1 m, headed with 60 cm of empty space. The latter space was occupied by excess water during runs. The water flow through the column was gravity assisted (speed of water through the column of sand was 0.15 meters per hour) with an outflow of water regulated at a speed of 4 L/h by a valve. An overflow pipe was used to maintain a constant water level above the sand column. The sand ratio consisted of different grain diameters of which 10% was > 1mm, 10% < 0.2mm, and 80% between 0.2mm and 1mm. This composition is considered the most effective to encourage the interaction between microorganisms and water, and allow the water to flow through the sand without being mechanically stopped (Pettitt, 2005). This experimental design determined half of the irrigation freshwater to be pumped to the top of the channels of the NFT-Type system, and the other half sent to the top of the column of the sand filter for biofiltration. Additional samples of sand were collected for molecular tests. These were collected via ‘sampling ports’ installed in the column (Fig. 1) at different layers of the SSF column: the top (1 cm), the middle (50 cm) and the bottom layer (80 cm).

### Dry weight of plants

Analysis of variance (ANOVA) was carried out on the dry weight of tomato plants. This allowed the comparison between the dry weight of the plants that were grown with recycled irrigation freshwater filtered with the slow sand filter (treatment plants, ssf-plants), and those grown with non-filtered recycled irrigation freshwater (control plants, co-plants).

### DNA isolation

For each DNA extraction three technical replicates were produced. When extracting from water, a negative control of sterile water was processed in parallel with samples of recycled irrigation freshwater to check for contamination. Genomic DNA was extracted from 50 ml of irrigation freshwater filtered onto a 0.2 μm Whatman nylon filter of 25 mm diameter (Horakova *et al*., 2008). After filtration, the 0.2 μm Whatman filter was placed in a microtube, with 0.5 g of glass beads (Sigma-Aldrich, Gillingham, UK) and 700 μl of extraction buffer. Each tube was shaken for 3 min in a Fastprep (QBiogene, Cambridge, UK) for cells disruption, for 4 cycles of 45 seconds at 6.5 m s^-1^. After fastprep, the supernatant was recovered. When extracting DNA from sand, 500 mg of sand were placed directly into the microtube with glass beads and extraction buffer, skipping the step of filtration with 0.2 μm Whatman filter.

To the recovered supernatant, 0.1% (w/v) of sodium dodecyl sulphate (SDS) were added, and the sample was homogenised and placed on ice for 10 min. Seven hundred μl of phenol:chloroform:isoamyl alcohol (25:24:1) were added, followed by centrifugation at 5 000 × g for 15 min. The aqueous phase was recovered, and the DNA was precipitated with 1.5 volume of isopropanol, and 0.1 volume of 0.5 M NaCl. The mixture was incubated overnight at -20°C. After incubation, a centrifugation step was carried out at 10 000 × g for 10 min, and the pellet was washed twice with 300 μl of 70% ethanol. DNA was finally eluted in 30 μl of sterile water. The DNA was further purified by polyvinylpyrrolidone (PVPP) cleanup method (Menking *et al*., 1999). One hundred mg of PVPP were added to a sterile spin column in a 1.5 ml tube. The column was washed twice with 300 μl of sterile water and centrifuged at 1 000 × g for 2 min. After washing, eluted DNA was added to the column and centrifuged at 1 000 × g for 2 min. DNA concentration was evaluated by 1.5% agarose gel electrophoresis, and by NanoDrop (ThermoScientific, Wilmington, USA).

### PCR amplification of ribosomal DNA

Isolated DNA was PCR amplified with the primer pairs for the portions 16S (Muyzer *et al*., 1993; Muyzer *et al*., 1995), and the 23S (Anthony *et al*., 2000) of the rRNA (Table 1). Amplifications were performed in a PTC200 thermocycler (MJ Research, St. Bruno, Quebec, Canada) in 25 μl reactions containing 1 μl of genomic DNA, 12.5 μl of 2X PCR MangoMix (Bioline, London, UK), 0.5 pmol of each primer and 10.5 μl of sterile distilled water. The following cycles were used for the DNA amplification: 1 cycle at 94°C for 2 min followed by 30 cycles of 94°C for 30 s, annealing (Table 1) for 1 min and 72°C for 2 min and 30 s, and a final extension step of 72°C for 10 min. Amplified DNA was verified by gel electrophoresis of aliquots of PCR mixtures (4 μl) in 1.5% of agarose in 1X TBE buffer and ethidium bromide (0.5 μg/ml).

**Table 1.**
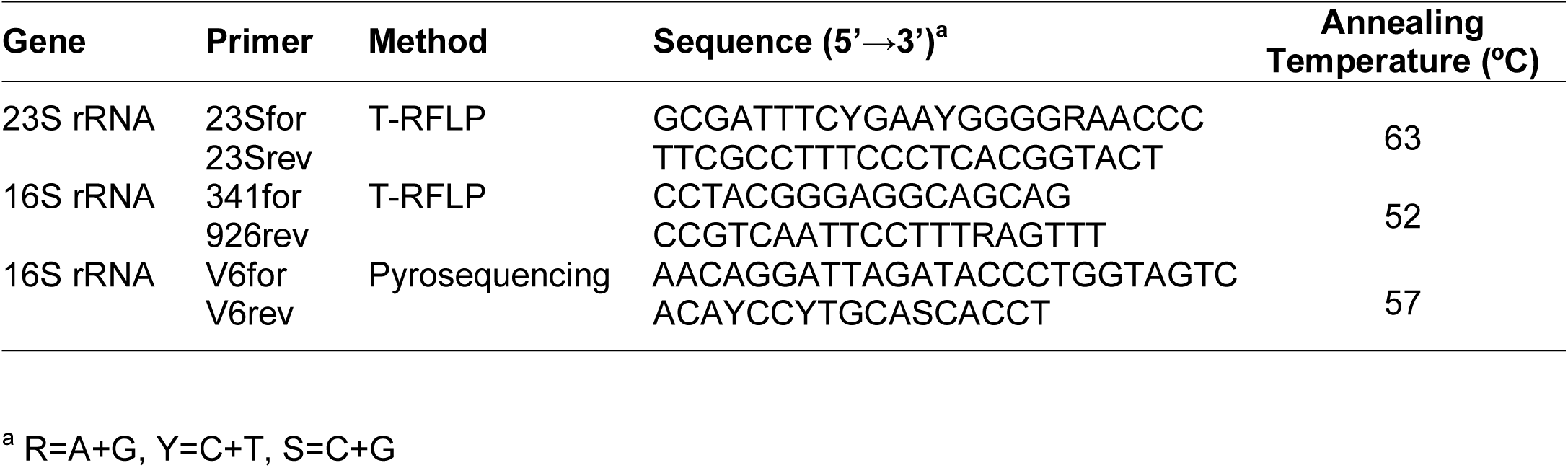
Primer pairs used in this study for the amplification of ribosomal DNA for T-RFLP and pyrosequencing

When PCR reactions were performed for T-RFLP, the reverse primer was fluorescently labelled with a Beckman dye at the 5’-end: for 16S rRNA, the primer 926rev was labelled with D3 (D3-926rev), and for 23S rRNA the primer 23Srev was labelled with D4 (D4-23Srev). All the fluorescent dyes are Beckman dyes provided by Sigma Proligo (Sigma-Aldrich). PCR products for T-RFLP were digested with two restriction enzymes (described below) to produce Terminal-Restriction Fragments (T-RFs) for each amplicon generated by PCR amplification. The combination of pairs of T-RFs was used for the identification of microorganisms.

### Restriction digestion

Ten μl of PCR product were digested in two separate reactions for each of the two restriction enzymes *Mse*I and *Hae*III (New England Biolabs, Hitchin, UK) in a 20 μl reaction volume containing 1U of restriction enzyme according to the manufacturer’s protocols. The digestion mix was incubated at 37°C for 2 hours, followed by enzyme denaturation by heating at 80°C for 20 min. Digestion products were verified by gel electrophoresis of aliquots of the digestion mixture (4 μl) in 2.5% of agarose in 1X TBE buffer and ethidium bromide (0.5 μg/ml).

### Terminal Restriction Fragment Length Polymorphism (T-RFLP)

T-RFLPs were determined by electrophoresis with a CEQ™ 8000, Genetic Analyzer System (Beckman Coulter, High Wycombe, UK). The product of restriction analysis was loaded into a 96 well plate with each well containing 38.5 μl of sample loading solution (GenomeLab, Beckman Coulter, High Wycombe, UK) and 0.5 μl of size standard-600 (GenomeLab, Beckman Coulter). Samples were covered with mineral oil and separated on the CEQ™ 8000. Digestions of 16S and 23S rRNA PCR amplicons were run in the same reaction because they were tagged with different dyes, D3 and D4 respectively, each of which is read at a different wavelength; to each well, 1.5 μl of each digestion product was loaded.

A quartic polynomial model was run for size standard calibration to improve correlation between expected and actual sizes (McEniry *et al*., 2008), particularly for fragments in the range 400-600bp (Brodie *et al*., 2002). T-RFs that differed by <0.5 bp in size between replicated profiles were considered identical and only T-RFs that occurred in at least two of the three replicates were included in the analyses (Dunbar *et al*., 2001). T-RFLP datasets were normalised dividing each peak height value by the sum of all peak height values in the correspondent profile (Hartmann and Widmer, 2008). Analysis of similarities (ANOSIM) was carried out on quality filtered T-RFLP datasets, with Bray-Curtis dissimilarity used to compute distance matrices of correspondence between samples. The ordination method non-metric multidimensional scaling (nMDS) was carried out on distance matrices with the software PAST (http://folk.uio.no/ohammer/past/), with a Shepard plot and a stress value reported for each plot.

Putative phylogenetic identities of T-RFs were assigned using a database obtained from the collection of known sequences from NCBI (http://www.ncbi.nlm.nih.gov). The DNA sequences were digested *in silico* with the software pDRAW32 (AcaClone Software, http://www.acaclone.com/), which identified all the restriction sites that were present in the DNA sequences. The database provided pairs of expected gene-enzyme combinations specific for each group of microorganisms, which were ultimately compared with experimental T-RFLP datasets.

### Purification of PCR products

PCR products, when used as inserts for cloning reactions, were purified with GenElute™ PCR Clean-Up Kit (Sigma-Aldrich) according to the manufacturer’s protocol. After elution in water, the concentration of PCR product was calculated with NanoDrop.

### Ligation and cloning

Purified ribosomal DNA amplicons were used as inserts for the ligation into pGEM®-T Easy Vector (Promega, Madison, WI USA). The ligation reaction was carried out in 10 μl volumes with 3U of T4 DNA ligase, 5 μl of 2X ligation buffer, 1μl of pGEM®-T Easy (50 ng/μl) and 22.5 ng of purified PCR product, and incubated overnight at 4°C. Promega *Escherichia coli* JM109 cells were transformed according to the manufacturer’s protocol.

Transformed cells were incubated at 37°C in Petri dishes with Luria Bertani (LB) medium, containing agar 15 g/L, IPTG (Isopropyl β-D-1-thiogalactopyranoside) 0.05 mM, X-GAL (5-bromo-4-chloro-3-indolyl-betagalactoside) 80 μg/ml and Ampicillin 100 μg/ml (Sambrook *et al*., 1989). White colonies (potential positive clones) were selected and screened by colony PCR with vector-targeted PCR: M13for (5’-GTAAAACGACGGCCAGT-3’) and M13rev (5’-CAGGAAACAGCTATGAC-3’) were used in 25 μl of PCR reaction in the same conditions as above, with an annealing temperature of 56°C. Amplified DNA was verified by 1.5% agarose electrophoresis. Positive clones from the colony PCR were checked by restriction analysis with the two restriction enzymes *Mse*I and *Hae*III. Clones displaying different banding patterns were selected for sequencing with a Beckman CEQ™8000 automated sequencer.

### Pyrosequencing of the V6 region of 16S rRNA

Pyrosequencing reads were obtained from PCR amplicons of the V6 hyper variable region of the 16S rRNA (Table 1) using a Roche 454 pyrosequencer (Roche, Basel, Switzerland) available at FERA (Food and Environment Research Agency, York, UK).

Amplifications were performed in a GeneAmp PCR system 9700 (Life Technologies Ltd, Paisley, UK) in 25 μl reaction volume containing 1 μl of genomic DNA, 5 μl of 5X KAPAHiFi fidelity buffer (KAPABIOSYSTEMS, Boston, MA USA), 0.75 μl of a mix of dNTPs 10 mM each, 0.5 pmol of each primer, 0.5 μl KAPAHiFi™ HotStart DNA polymerase and sterile distilled water. The following cycling conditions were used: 1 cycle at 95°C for 4 min followed by 35 cycles of 98°C for 20 sec, annealing at 57°C for 15 sec and 72°C for 30 sec, with a final extension step of 72°C for 5 min. Amplified DNA was verified by gel electrophoresis of aliquots of the PCR mixtures (4 μl) in 1.5% of agarose in 1X TBE buffer and ethidium bromide (0.5 μg/ml). The forward primer contained the adapter forward (5’-CGTATCGCCTCCCTCGCGCCATCAG-3’), and the reverse primer the adapter reverse (5’-CTATGCGCCTTGCCAGCCCGCTCAG-3’). These two adapters were used in the library preparation step of pyrosequencing to create a bond between single stranded amplicons and glass beads. Furthermore, the forward primer was provided of a specific 10 nucleotide (-nt) multiple identifier (MID) that was added to the 3’-end of the adapter and to the 5’-end of the forward primer. A set of 4 different MIDs (MID1 to 4 according to manufacturer’s indications) was used to combine different samples in the same reaction mixture, and retrieve the original sample composition at the end of the pyrosequencing reaction. PCR products of an approximate length of 280-nt were quantified with PicoGreen (Life Technologies) followed by quality analyses performed according to the manufacturer’s instructions. Equal amounts of samples were mixed in groups of 4 (containing the 4 different MIDs) and run overnight for the pyrosequencing reaction.

### Pyrosequencing data analysis

Raw pyrosequencing data were further analysed after the quality filtering provided by the 454 pyrosequencer. Samples composition was investigated through diversity measures, while the phylogenetic identity of the DNA reads was evaluated with a) RDP classifier (Cole *et al*., 2009) and b) BLAST searches, followed by MEGAN (Huson et al., 2007).

The software mothur (Schloss *et al*., 2009) and the RDP pipeline (Cole *et al*., 2009) were used in combination for the trimming, alignment and clustering of the pyrosequencing reads. The statistical and graphics software package R (www.r-project.org) was used to generate rarefaction curves, and the software EstimateS (Colwell, 2006) to produce diversity indices. In addition, diversity measures were used to compare the two datasets obtained with T-RFLP and pyrosequencing (Table 5).

Quality filtered sequencing reads were trimmed according to published recommendations (Huse *et al*., 2007) using the RDP pyrosequencing pipeline (Cole *et al*., 2009), with reads shorter than 50bp and containing ambiguous bases excluded from the analysis. Pyrosequencing reads were aligned with Infernal (Nawrocki and Eddy, 2007), followed by complete linkage clustering of the RDP pipeline.

The Naïve Bayesian classifier RDP-classifier (Wang *et al*., 2007) provided rapid phylogenetic assignment for taxonomic classification. This method was used to compare the DNA reads obtained from filtered recycled irrigation freshwater (Water *sff*), the control recycled irrigation freshwater (Water co), and the sand of the top layer of the SSF (Sand *top*) throughout the 4 time points.

The analysis of the time point ‘28 days’ was implemented with a BLASTn analysis followed by the phylogenetic analysis with the software MEGAN (Huson *et al*., 2007). MEGAN provides hierarchical tree construction based on BLAST searches (Altschul *et al*., 1990), assigning the 16S rRNA gene reads to NCBI taxonomy. MEGAN also calculates relative abundances of pyrosequencing reads among samples. BLAST searches give a more accurate analysis of query sequences as compared to the RDP classifier; however, the production of the BLASTn file (input file for MEGAN) is highly time consuming and requires powerful processing units. For this reason, BLASTn files were produced for the ‘28 days’ time point only, with the assumption that this time point would be the most informative of the 4.

### Diversity indices and richness estimators

The diversity of bacteria in recycled irrigation freshwater was estimated to compare the molecular methods T-RFLP and pyrosequencing (Table 5). The richness estimators Chao1 (Chao and Bunge, 2002) and ACE (Chao *et al*., 2006) were used for the analysis of pyrosequencing data.

## RESULTS

### Plant growth

The dry weight of the plants *Solanum lycopersicum* cv. Alicante, grown in the experimental NFT-Type systems were used to assess the effect of slow sand filtration on plant growth. ANOVA (Figure 2) showed that plant growth was significantly affected by the filtration with the slow sand filter (F_1,18_ = 4.78, p = 0.036). Plants grown with filtered recycled irrigation freshwater (ssf-plants) had a larger biomass than those grown with control recycled irrigation freshwater (co-plants).

**Figure 2.**
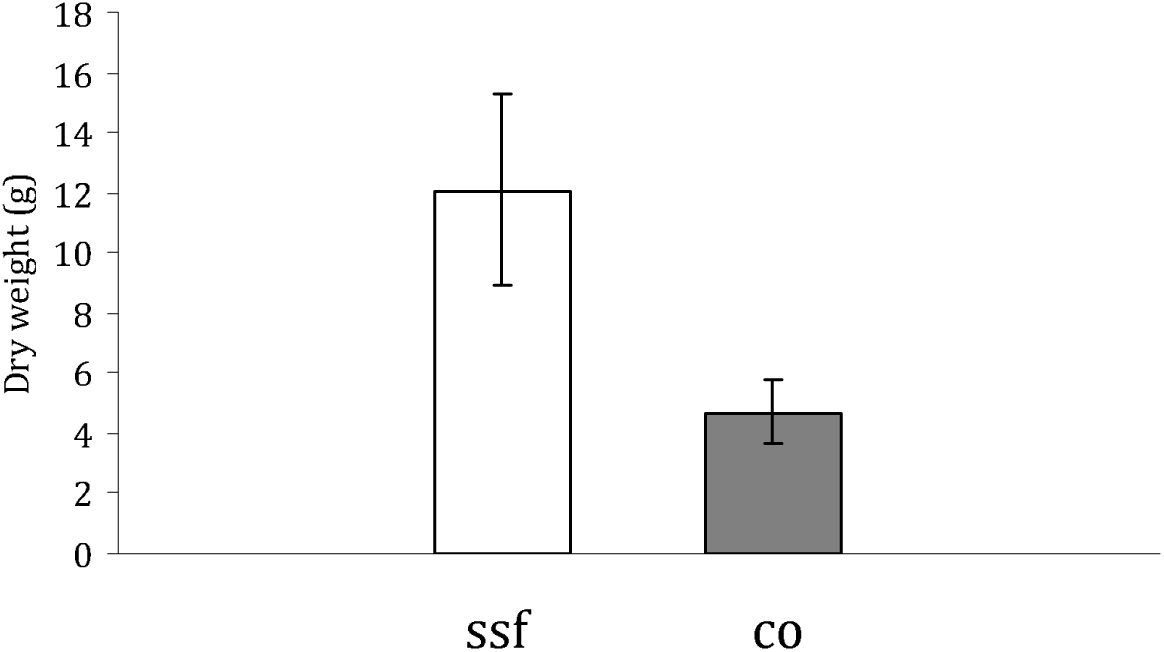
Dry weight of tomato plants grown in the NFT-type experimental system. Error bars indicate S.E.M.

### Bacterial Community Composition (BCC) with T-RFLP

ANOSIM was performed on T-RFLP datasets (Table 2) to test the effect of the slow sand filtration (*H_0_* ssf) and time (*H_0_* time) on the BCC of recycled irrigation freshwater. Slow sand filtration and time had a significant effect on the BCC of recycled irrigation freshwater. Datasets were analysed to test the null hypothesis that i) there were no differences between datasets over the 4 time points (*H_0_* time); and ii) there were no differences in the BCC between filtered recycled irrigation freshwater (Water *ssf*) and control recycled irrigation freshwater (Water *co*) (*H_0_* ssf). The results from 16S and 23S rRNA genes showed R-values between 0.8 and 1 with all gene-enzyme combinations, and were supported by low probability values (all <0.0001).

**Table 2.** Observed two-way ANOSIM test values and probabilities of null hypotheses (*H_0_*) tests obtained comparing T-RFLP datasets of 16S and 23S rRNA genes with the enzymes *Mse*I and *Hae*III. *H_0_* was tested between time points (*H_0_* time) and water sample coming from recycled irrigation water treated with the slow sand filter and control water (*H_0_* ssf)

| NFT | Gene | *MseI* |  | *HaeIII* |  |
| --- | --- | --- | --- | --- | --- |
|  |  | R-value | Probability | R-value | Probability |
| NFT1 | | $H_0$ time | | | |
|  | 16S | 0.81778 | < 0.0001 | 0.93519 | < 0.0001 |
|  | 23S | 0.99537 | < 0.0001 | 0.82469 | < 0.0001 |
| | | $H_0$ ssf | | | |
|  | 16S | 0.85417 | < 0.0001 | 1 | < 0.0001 |
|  | 23S | 0.87963 | < 0.0001 | 1 | < 0.0001 |
| NFT2 | | $H_0$ time | | | |
|  | 16S | 0.98611 | < 0.0001 | 0.96759 | < 0.0001 |
|  | 23S | 0.89198 | < 0.0001 | 0.94599 | < 0.0001 |
| | | $H_0$ ssf | | | |
|  | 16S | 0.98148 | < 0.0001 | 0.99074 | < 0.0001 |
|  | 23S | 0.87037 | < 0.0001 | 0.92593 | < 0.0001 |
| NFT3 | | $H_0$ time | | | |
|  | 16S | 0.77006 | < 0.0001 | 0.97840 | < 0.0001 |
|  | 23S | 1 | < 0.0001 | 1 | < 0.0001 |
| | | $H_0$ ssf | | | |
|  | 16S | 0.83333 | < 0.0001 | 0.97222 | < 0.0001 |
|  | 23S | 1 | < 0.0001 | 1 | < 0.0001 |

Multivariate statistical tests of T-RFLP datasets with nMDS showed that samples of Water *ssf* occupied the same part of the plot starting at 14 days of filtration (Fig. 3). This was particularly clear with the gene-enzyme combination 23S-*MseI*: the nMDS plot shows that after 7 days, Water *ssf* and Water *co* (black spots and red crosses respectively) grouped together on the left hand side of the plot. For the other time points, samples of Water *co* plotted in the top right hand side, while Water *ssf* grouped together and seemed to stabilise in the same position of the plot after 14, 21 and 28 days.

**Figure 3.**
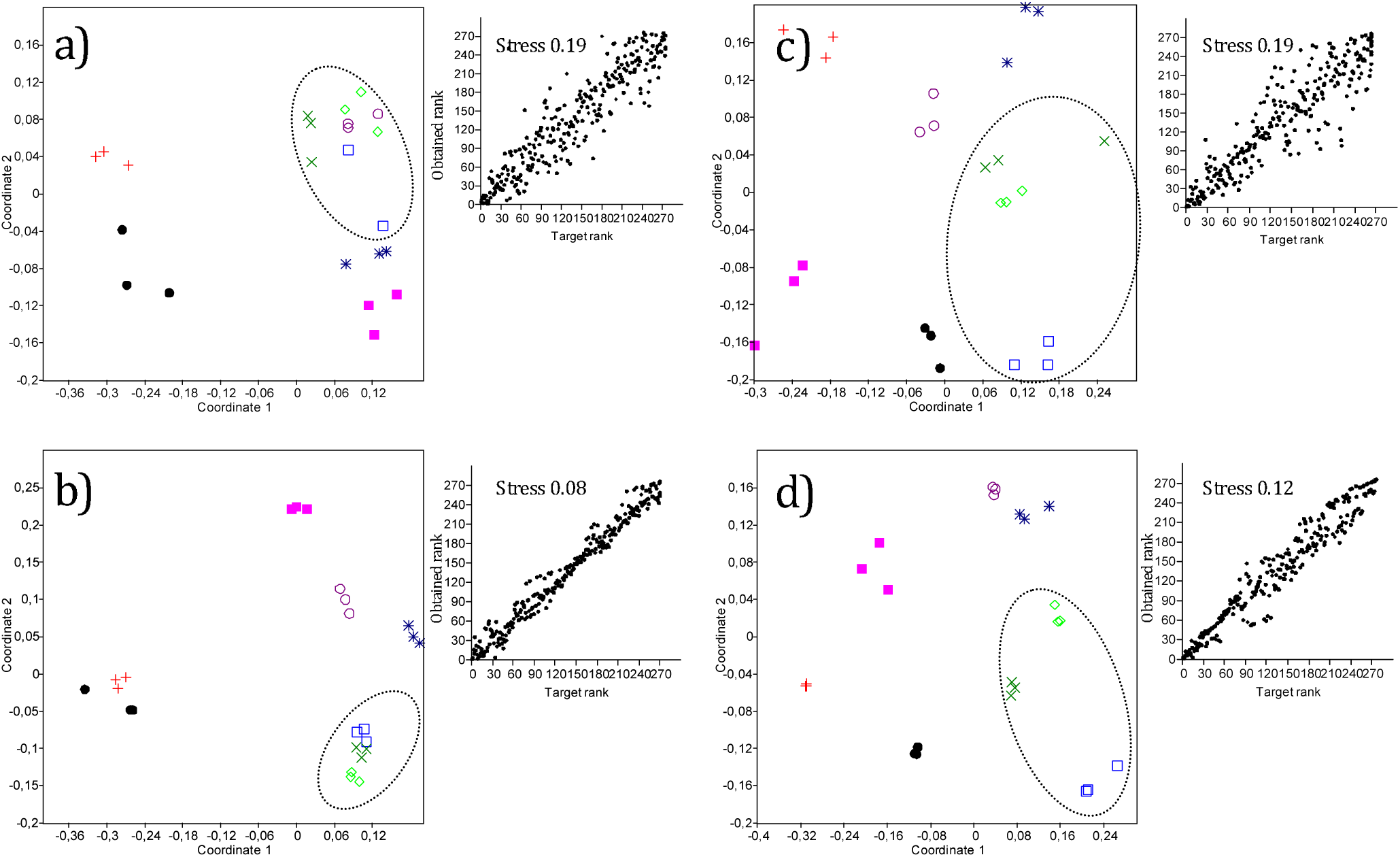
Non-metric multidimensional scaling of microorganisms inhabiting the recycled irrigation water of the experimental NFT system, using T-RFLP profiles of 16S rRNA (a, c) and 23S rRNA (b, d). Each shape represents a different time point and a different water sample of filtered (Water *ssf*) and non filtered (Water *co*) recycled irrigation water: black dots 7ssf; red crosses are 7co; blue squares 14ssf; pink filled squares 14co; green crosses 21ssf; purple circles 21co; green diamonds 28ssf; and blue stars 28co. Elliptic shapes group samples of filtered (*ssf*) recycled irrigation water after 14, 21 and 28 days. Each rRNA gene is represented by two profiles obtained with two different restriction enzymes: 16S *Mse*I (a) and *Hae*III(c), 23S MseI (b) and HaeIII(d). Each nMDS plot is provided with correspondent stress value and regression analysis.

### Pyrosequencing of the V6 region of 16S rRNA gene

Pyrosequencing analysis was performed on the hypervariable V6 region of the 16S rRNA. A total of 332 608 DNA reads were obtained by 454 pyrosequencing (Table 3). Of those, 291 442 (87.6%) passed the quality filter check, with an average length of 211 bp. DNA reads were further analysed with the RDP pyrosequencing pipeline and mothur.

**Table 3.** Data summary of total reads from pyrosequencing data. “Trimmed reads” represent the number of DNA sequences longer than 50bp that were kept after quality filtering. “Unique reads” represent the number of distinct sequences within a set of Trimmed reads

| Sample ID | Total reads | Trimmed reads | Unique reads |
| --- | --- | --- | --- |
| 7ssf | 25005 | 22951 | 11237 |
| 14ssf | 31852 | 28133 | 15083 |
| 21ssf | 34518 | 29400 | 15282 |
| 28ssf | 23580 | 21125 | 11525 |
| 7co | 21080 | 16967 | 7367 |
| 14co | 58995 | 52248 | 23295 |
| 21co | 21107 | 19845 | 8872 |
| 28co | 8808 | 6740 | 3803 |
| 7top | 22842 | 21191 | 10697 |
| 14top | 42472 | 38961 | 19473 |
| 21top | 26581 | 22842 | 11723 |
| 28top | 15768 | 11039 | 6131 |
| $\Sigma$ | 332608 | 291442 | 144488 |

For the description of the communities, trimmed sequences were clustered into groups of defined sequence variation that ranged from unique sequences (no variation) to 10% differences by mothur. These clusters were then used to plot OTUs versus the number of tags for generating rarefaction curves (data not shown) and for obtaining richness estimators such as the abundance-coverage estimator (ACE) and Chao1. These indices showed that the species diversity estimation was of the third order of magnitude. At 97% similarity, Chao1 predicted that the maximum number of OTUs was between 11 016 and 15 100 for Water *ssf*, 4 530 and 17 369 for Water *co* and 7 271 to 16 126 for sand of the top layer (Sand *top*) (Table 4).

**Table 4.** Similarity-based OTUs and species richness estimators

| Sample ID | Trimmed reads | Cluster distance |  |  |  |  |  |  |  |  |
| --- | --- | --- | --- | --- | --- | --- | --- | --- | --- | --- |
|  |  | 0.03 |  |  | 0.05 |  |  | 0.1 |  |  |
|  |  | OTUs | ACE | Chao1 | OTUs | ACE | Chao1 | OTUs | ACE | Chao1 |
| 7ssf | 22951 | 4942 | 18053 | 11016 | 3848 | 11713 | 7819 | 2730 | 6831 | 4929 |
| 14ssf | 28133 | 6344 | 22750 | 13970 | 4900 | 14660 | 9875 | 3434 | 8449 | 6115 |
| 21ssf | 29400 | 6634 | 23351 | 15100 | 5126 | 16287 | 10828 | 3602 | 8582 | 6221 |
| 28ssf | 21125 | 5684 | 22455 | 13288 | 4642 | 16133 | 9893 | 3385 | 9673 | 6549 |
| 7co | 16967 | 3107 | 10495 | 6868 | 2378 | 7269 | 4936 | 1648 | 4469 | 3165 |
| 14co | 52248 | 8468 | 26004 | 17369 | 6211 | 16225 | 11847 | 3954 | 8314 | 6801 |
| 21co | 19845 | 3567 | 10942 | 7436 | 2661 | 6893 | 4953 | 1738 | 3729 | 3053 |
| 28co | 6740 | 1883 | 7916 | 4530 | 1522 | 5340 | 3397 | 1102 | 3448 | 2258 |
| 7top | 21191 | 4928 | 18547 | 11588 | 3768 | 12325 | 8001 | 2564 | 7089 | 4875 |
| 14top | 38961 | 7507 | 24877 | 16126 | 5604 | 15738 | 10928 | 3624 | 7872 | 5976 |
| 21top | 22842 | 5244 | 17773 | 11664 | 4087 | 12364 | 8425 | 2820 | 7378 | 5257 |
| 28top | 11039 | 3147 | 11409 | 7271 | 2518 | 8312 | 5440 | 1825 | 5921 | 3757 |

ACE and Chao1 constantly estimated a number of OTUs higher than those that were effectively obtained, suggesting that sampling represented a limitation for the identification of the total number of species.

### Bayesian hierarchical identification of pyrosequencing reads

Pyrosequencing reads of recycled irrigation freshwater and the sand of the top layer of the SSF were phylogenetically assigned with the Naïve Bayesian classifier of the RDP-classifier (Fig. 4). Major changes were observed for the classes of *Bacilli, Alpha-* and *Gammaproteobacteria* and *Nitrospira*. The latter taxon was more abundant in Water *ssf* and Sand *top*, while only a minor number of reads were detected in Water *co. Alphaproteobacteria* decreased in relative abundance throughout the 4 time points. *Gammaproteobacteria* were less abundant in Water *ssf* after 28 days as compared to Water *co. Bacilli* increased their relative abundance in three of the four time points. Only after 21 days, *Bacilli* were more abundant in the Water *co* as compared to Water *ssf*.

**Figure 4.**
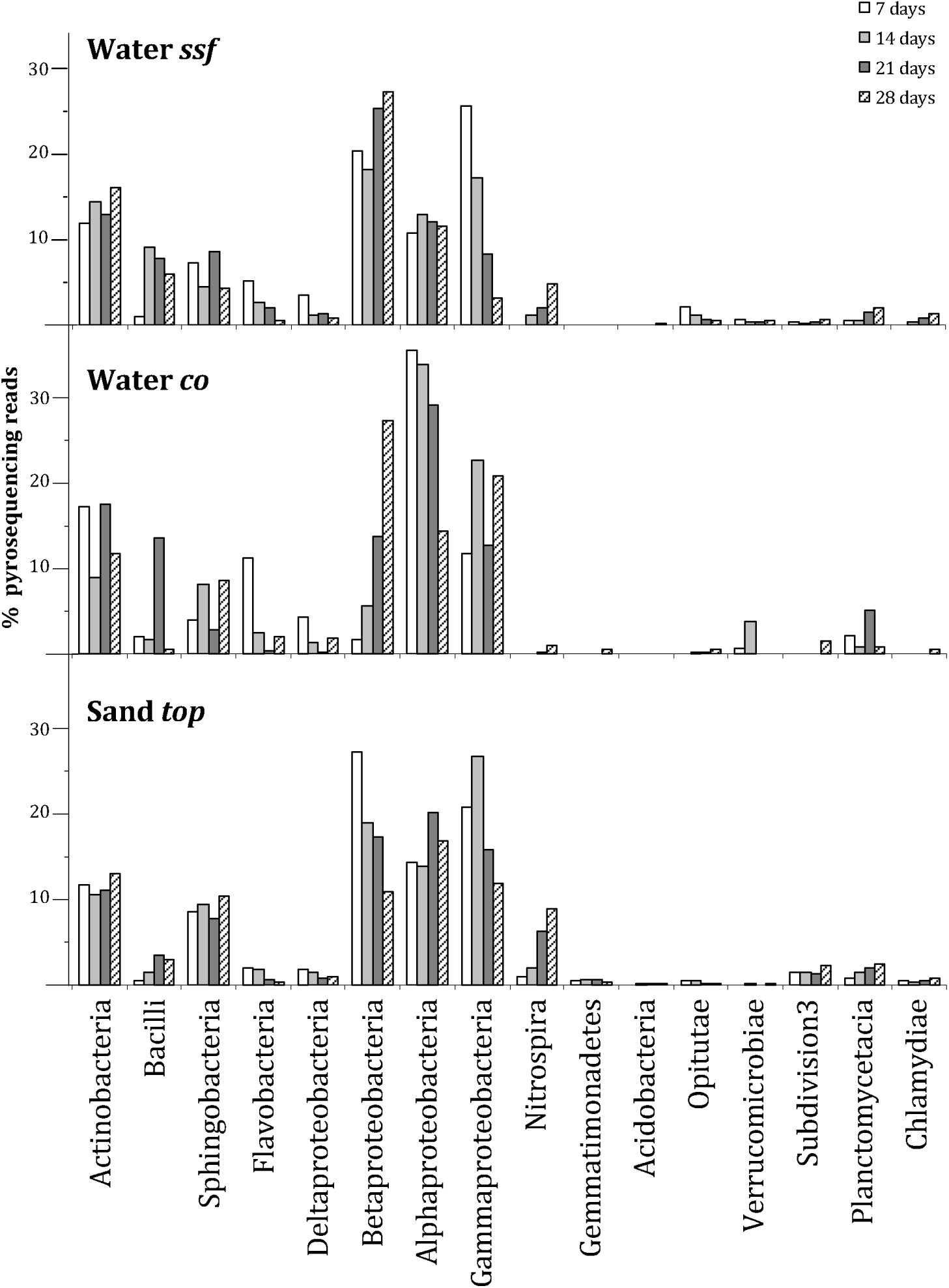
Major classes of bacteria identified with the pyrosequencing of the V6 region of bacterial 16S rRNA. Plots show relative abundance (percentage) of the reads obtained at the 4 time points (7, 14, 21 and 28 days) for a) filtered recycled irrigation water (Water *ssf*), b) control recycled irrigation water (Water *co*) and c) top layer of the slow sand filter (Sand *top*).

### Phylogenetic identification of pyrosequencing reads using MEGAN

To provide a higher resolution of phylogenetic assignment beyond the class level, recycled irrigation freshwater and sand of the top layer of the SSF were analysed with BLASTn and MEGAN (Fig 5). Slow sand filtration increased the number of taxa in the recycled irrigation freshwater from 26 of Water *co* to 45 of Water *ssf*. The three investigated samples (Water *ssf* after 28 days-28 *ssf;* Water *co* after 28 days-28 *co;* and Sand *top* after 28 days-28 *top*) shared 17 genera, with only 6 shared by Water *ssf* and Water *co*. Overall, the most abundant phyla were *Actinobacteria, Bacteroidetes, Firmicutes* and *Proteobacteria*.

**Figure 5.**
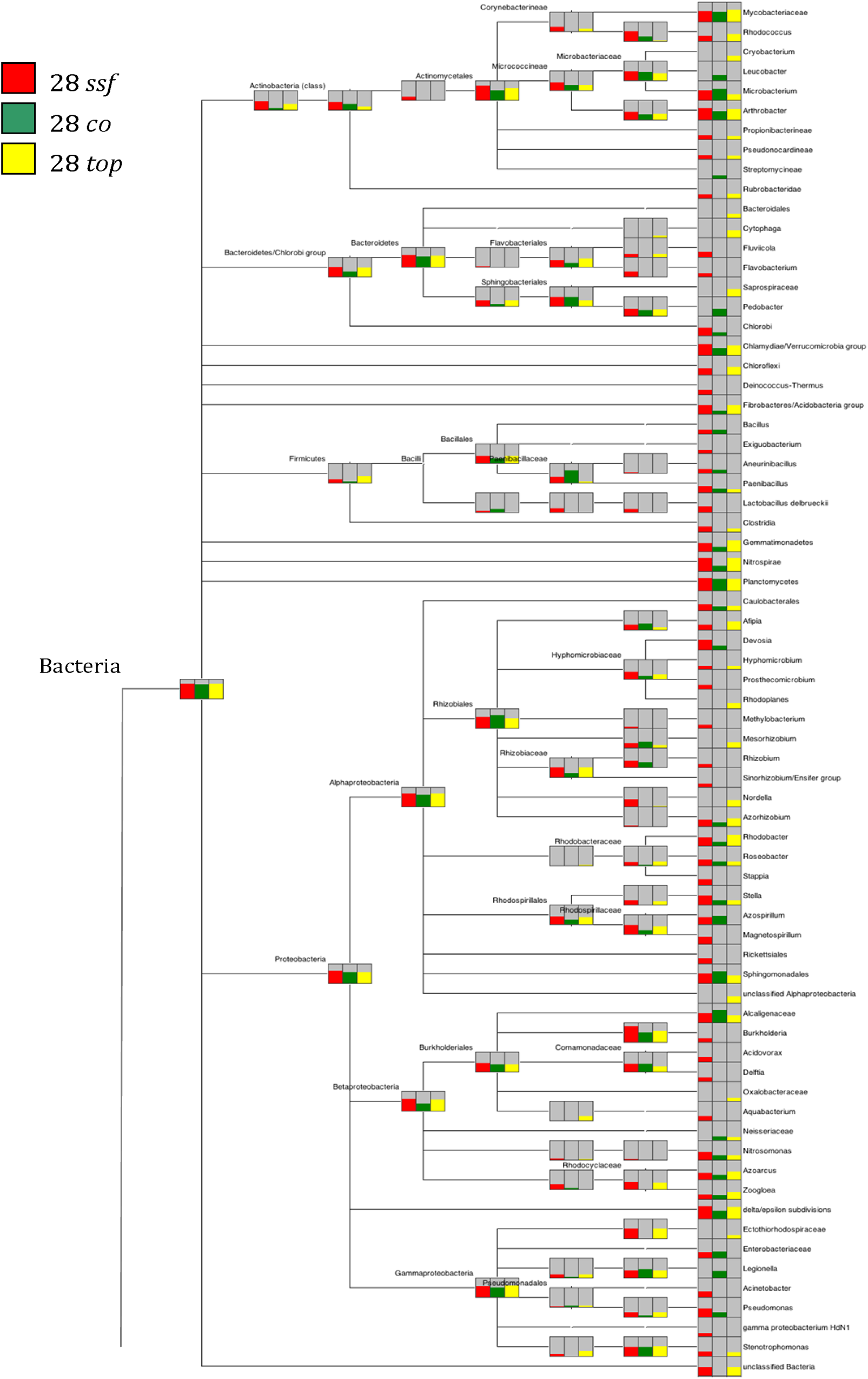
V6 amplicon sequences from the three groups of samples assigned with BLAST and MEGAN. Coloured bars display the relative abundance for each taxon of bacteria.

Several groups of bacteria were exclusively present in Water *ssf* (*28 ssf*). These taxa included *Fluviicola* and *Flavobacterium* of the class of *Bacteroidetes, Deinococcus* of the class of *Deinococci, Exiguobacterium* and *Lactobacillus* of the class of *Bacilli, Prosthecomicrobium, Methylobacterium, Rhizobium, Sinorhizobium, Stappia, Magnetospirillum* and unclassified *Rickettsiales* of the class of *Alphaproteobacteria, Burkholderia, Acidovorax, Delftia* and *Aquabacterium* of the class of *Betaproteobacteria*, and finally *Acinetobacter* and unclassified *Gammaproteobacteria*.

Other groups of bacteria increased their relative abundance after 28 days of filtration with the SSF. These included unclassified *Gemmatimonadetes, Nitrospira* of the phylum *Nitrospira, Azorhizobium, Rhodobacter, Roseobacter* and *Stella* of the class of *Alphaproteobacteria, Nitrosomonas* and *Azoarcus* of the class of *Betaproteobacteria, Bdellovibrio* and *Geobacter* of the class of *Deltaproteobacteria*, and finally *Pseudomonas* of the class of *Gammaproteobacteria*. Interestingly, two microorganisms responsible for two different step of the oxidation of nitrogen were enhanced by the slow sand filtration, such as *Nitrosomonas* and *Nitrospira*. In particular, the latter showed a large increase in number of reads between Water *ssf* and Water *co*, with 1749 reads against 18.

Other microorganisms showed a reduction of relative abundance between Water *ssf* and Water *co*. These included *Microbacterium* of the class of *Actinobacteria, Sphingomonas* and *Azospirillum* of the class of *Alphaproteobacteria*, unclassified *Alcaligenaceae* of the class of *Betaproteobacteria* and unclassified *Enterobacteriaceae* of the class of *Gammaproteobacteria*.

Finally, some organisms did not appear in Water *ssf*, while did in Water *co*. These included, *Leucobacter* and *Streptomyces* of the class of *Actinobacteria, Pedobacter* of the class of *Sphingobacteria*, unidentified *Neisseriaceae* of the class of *Betaproteobacteria*, and *Legionella* of the class of *Gammaproteobacteria*.

### Comparison of pyrosequencing and T-RFLP

T-RFLP and pyrosequencing results showed some degrees of similarity. T-RFLP provided a rapid overview of changes in dominant members of the community and, when implemented with clone libraries, their phylogenetic identification. However, pyrosequencing data provided in depth information about changes in the relative amount of thousands of OTUs, which were, at least, of 2 orders of magnitude higher than T-RFLP.

T-RFLP underestimated the diversity of the systems expressed by Shannon and Simpson indices (Table 5). These two indicators were always lower for T-RFLP data. Nonetheless, T-RFLP provided reliable information about major groups of bacteria and their patterns of relative abundances. 16S rRNA T-RFLP detected the increase of *Pseudomonas* sp., *Cellvibrio* sp. and *Nitrosospira* sp. and the decrease of *Novosphingobium* sp. in Water *ssf* as compared to Water *co* (Table 6). When T-RFLP was performed on 23S rRNA amplicons, an increase was detected for *Azoarcus aromaticum, Paenibacillus* sp. and *Ochrobactrum intermedium*, while a decrease was detected for *Sphingomonas* sp.

**Table 5.** Number of OTUs and diversity indices obtained from T-RFLP and pyrosequencing of the 16S region of rRNA

| Sample ID | T-RFLP |  |  |  |  |  | Pyrosequencing cluster distance |  |  |  |  |  |  |  |  |
| --- | --- | --- | --- | --- | --- | --- | --- | --- | --- | --- | --- | --- | --- | --- | --- |
|  | 16S-*Mse*I |  |  | 16S-*Hae*III |  |  | 0.03 |  |  | 0.05 |  |  | 0.1 |  |  |
|  | OTUs | *H'*<sup>a</sup> | 1-D | OTUs | *H'* | 1-D | OTUs | *H'* | 1-D | OTUs | *H'* | 1-D | OTUs | *H'* | 1-D |
| 7ssf | 4 | 1.26 | 0.66 | 4 | 1.42 | 0.71 | 4942 | 5.43 | 0.98 | 3848 | 5.21 | 0.98 | 2730 | 4.78 | 0.97 |
| 14ssf | 5 | 0.96 | 0.42 | 3 | 1.06 | 0.64 | 6344 | 5.64 | 0.99 | 4900 | 5.41 | 0.99 | 3434 | 4.93 | 0.98 |
| 21ssf | 16 | 2.54 | 0.90 | 17 | 2.54 | 0.90 | 6634 | 5.52 | 0.98 | 5126 | 5.28 | 0.98 | 3602 | 4.83 | 0.97 |
| 28ssf | 13 | 2.20 | 0.84 | 13 | 2.12 | 0.82 | 5684 | 5.79 | 0.97 | 4642 | 5.56 | 0.97 | 3385 | 5.06 | 0.96 |
| 7co | 11 | 2.07 | 0.84 | 5 | 1.43 | 0.72 | 3107 | 4.55 | 0.97 | 2378 | 4.34 | 0.97 | 1648 | 3.94 | 0.96 |
| 14co | 6 | 1.63 | 0.77 | 24 | 2.89 | 0.93 | 8468 | 5.19 | 0.98 | 6211 | 5.00 | 0.98 | 3954 | 4.62 | 0.98 |
| 21co | 15 | 2.53 | 0.91 | 6 | 1.56 | 0.76 | 3567 | 4.87 | 0.97 | 2661 | 4.70 | 0.97 | 1738 | 4.24 | 0.96 |
| 28co | 16 | 2.50 | 0.90 | 18 | 2.61 | 0.91 | 1883 | 4.90 | 0.98 | 1522 | 4.67 | 0.98 | 1102 | 4.19 | 0.97 |
| 7top | 14 | 2.24 | 0.86 | 13 | 2.04 | 0.80 | 4928 | 5.50 | 0.98 | 3768 | 5.28 | 0.98 | 2564 | 4.82 | 0.98 |
| 14top | 11 | 2.06 | 0.83 | 21 | 2.65 | 0.90 | 7507 | 5.51 | 0.98 | 5604 | 5.27 | 0.97 | 3624 | 4.74 | 0.97 |
| 21top | 22 | 2.78 | 0.92 | 18 | 2.51 | 0.89 | 5244 | 5.73 | 0.96 | 4087 | 5.50 | 0.96 | 2820 | 4.93 | 0.94 |
| 28top | 28 | 3.10 | 0.95 | 18 | 2.62 | 0.91 | 3147 | 5.52 | 0.98 | 2518 | 5.30 | 0.98 | 1825 | 4.84 | 0.98 |

<sup>a</sup> *H'* = Shannon index; 1-D = Simpson's index

**Table 6.** Major microorganisms identified by T-RFLP and clone libraries. The column Relative abundance indicates the increase (arrows going upwards), or the decrease (arrows going downwards) of the relative amount of fluorescence in filtered water (Water *ssf*) as compared to control water (Water *co*)

| Gene | Genus/Species | T-RFs |  | Relative abundance |
| --- | --- | --- | --- | --- |
|  |  | *MseI* | *HaeIII* |  |
| 16S rRNA | *Cellvibrio* sp. | 589 | 509 | ↑* |
|  | *Nitrosospira* sp. | 588 | 588 | ↑* |
|  | *Novosphingobium* sp. | 564 | 484 | ↓ |
|  | *Pseudomonas* sp. | 374 | 31 (585) | ↑* |
| 23S rRNA | *Azoarcus aromaticum* | 372 | 312 | ↑* |
|  | *Paenibacillus* sp. | 422 | 169 | ↑ |
|  | *Ochrobactrum intermedium* | 515 | 201 | ↑ |
|  | *Sphingomonas* sp. | 337 | 202 | ↓ |

\* indicates that the difference of relative abundance between filtered water (Water *ssf*) and control water (Water *co*) was statistically significant

## DISCUSSION

A strong differentiation of the bacterial community composition (BCC) in recycled irrigation freshwater was triggered by the activity of the slow sand filter (SSF). The bioactive layer of the SSF, the schmutzdecke, affects the BCC of recycled irrigation freshwater. Specifically, phylogenetic affiliation of DNA reads revealed that the most affected classes of bacteria were *Bacilli, Alpha-* and *Gammaproteobacteria*, and *Nitrospira*. In addition, a larger diversity of bacteria was detected in recycled irrigation freshwater filtered with the SSF (Water *ssf*), as compared to control recycled irrigation freshwater (Water co): after 28 days of a NFT-type experiment, pyrosequencing data detected 45 taxa in Water *ssf*, against 26 in Water *co*. In addition, plant growth was enhanced when recycled irrigation freshwater was filtered with the SSF (Fig. 2). The activity of the schmutzdecke has been previously reported to be able to affect the BBC (Joupert and Pillay, 2008), but also to affect the abundance of common plant pathogens such as zoosporic fungi (Calvo-Bado *et al*., 2003; Calvo-Bado *et al*., 2006; Deniel *et al*., 2006). Further experiments were carried out to estimate whether the schmutzdecke needs adaption to be able to carry out its function in the SSF. Such tests confirmed that maturity is reached after 8 weeks (data not shown). The qualities of a SSF are promising for future applications of a combined system NFT-SSF: NFT guarantees a controlled closed environment for the growth of plants, while the SSF secures a microbiological balance of the system, which, in our experiments, has shown to enhance plant growth (Fig. 2).

Four main phyla of bacteria were detected as major components of the microflora of recycled irrigation freshwater. These included *Actinobacteria, Proteobacteria, Firmicutes Bacteroidetes*. High diversity was detected within these groups of bacteria, indicating that the nutrient solution of a hydroponic system consists of a rich microbiological environment (Berkelmann *et al*., 1994). Major taxa within the above phyla that were identified in this study have been previously described as common inhabitant of nutrient solutions. These included *Pseudomonadaceae, Xanthomonadaceae, Rhizobiaceae* and *Bacillaceae* (Berkelmann *et al*., 1994; Koohakan *et al*., 2004; Calvo-Bado *et al*., 2006). In addition, several species detected in this study have been previously reported as efficient biocontrol agent against the common plant pathogen *Fusarium oxysporum*. These included *Bacillus subtilis* (Baysal *et al*., 2008), *Bacillus megaterium* and *Burkolderia cepacia* (Omar *et al*., 2006),

The activity of the SSF determined a change in the BCC of recycled irrigation freshwater. This study showed that SSF influenced the relative abundance of a set of bacterial genera such as *Pseudomonas, Bacillus, Flavobacterium, Burkholderia* and *Azospirillum*. These bacteria have been previously described as plant growth promoting rhizobacteria (PGPR) (Glick, 1995; Lucy *et al*., 2004). Several reports have suggested that PGPR stimulate plant growth by facilitating the uptake of minerals into the plant (Kloepper *et al*., 1988). However, there is some controversy regarding the mechanisms that PGPR employ in the uptake of minerals (Bashan *et al*., 1990). Increased plant growth was observed when the nutrient solution was filtered with the SSF, after the period of ripening. However, no evidence has been previously reported of plant growth promotion by slow sand filtration. To support such findings, further work should be addressed at the isolation and deployment of specific consortia of microorganisms, in order to test their efficiency alone. Ideally, this should facilitate the formulation of recipes of consortia of microorganisms with which recycled nutrient solution in soilless hydroponic systems should be enriched. The presence, increased or exclusive, in Water *ssf* of bacteria such as *Bacteroidetes, Gemmatimonadetes, Nitrospira, Firmicutes, Alpha-, Beta-, Gamma-* and *Deltaproteobacteria*, was associated with an increased biomass in plants. However, it remains unclear whether physical-chemical interactions in the sand bed of the SSF actively contribute to the plant growth promoting effect. For instance, small scale sand filters could be used to prime nutrient solutions of recycled irrigation water, enabling the enrichment of the bacterial population, followed by an incubation period for the multiplication of the bacteria before employing the solution in hydroponic systems. The use of a slow sand filter represents a natural and inexpensive biological solution to enrich the bacterial population of recycled irrigation water, in a system where higher diversity reduces risks of colonization of single species (Stecher *et al*., 2010). Furthermore, these results increase scepticism on the use of disinfection methods that greatly reduce the microbial content such as UV disinfection (Zhang and Tu, 2000) and heat treatments (Runia *et al*., 1988). These findings suggest that a higher biodiversity of the BCC in the recycled irrigation freshwater has positive effects for the ecosystem, in which multiple interactions among species of several different phyla contribute to the efficiency of the NFT-Type experimental system.

The integration of fingerprinting and next generation sequencing provided the identification of microorganisms in recycled irrigation water. In this work, T-RFLP was developed for detecting microorganisms in water, using experimental tests to estimate the reliability and sensitivity of the method. These procedures showed that bacteria inoculated in sterile water are detected when their concentration is above the threshold of 10^3^ cells ml^-1^, and that an identical T-RFLP profile is obtained from overlaid T-RFLP profiles of single bacterial colonies, after DNA isolation/PCR/T-RFLP procedures, only when the concentration of cells in freshwater is at least of 10^6^ ml^-1^ (data not shown). At this concentration, T-RFLP is reliable and able to estimate relative amount of single species. T-RFLP has the advantage of being a rapid and reliable method that, at affordable prices, provides an overview of the system nonetheless. Another advantage of T-RFLP is that it provides datasets that can used to test complex hypotheses with multivariate statistics, which remains extremely useful when comparing different environments. In addition, this method showed complementarity with pyrosequencing results. Other authors (Jakobsson *et al*., 2010) have reported congruence between T-RFLP and pyrosequencing results. Our data supported such findings, showing that T-RFLP provides accurate information about dominant groups of microorganisms. Pyrosequencing, on the other hand, provides in depth information on thousands of operational taxonomic units (OTUs).

The analysis of pyrosequencing data produced useful species richness estimators that facilitated the ecosystem characterization. For example, Chao1 (Chao *et al*., 2006) was used to predict the species diversity of recycled irrigation water. This estimation, on average, showed that 50 ml contained around 10 000 OTUs at 97% similarity, going down to around 5 000 OTUs at 90% similarity. Other authors have previously estimated this diversity: Berkelmann *et al*. (1994) showed that the nutrient solution of a hydroponic system with rockwool as substrate, can contain up to 10^5^-10^6^ colony-forming units (CFU) ml^-1^ after 20 hours from planting tomato plants. Other attempts to estimate richness have mainly focused on other matrices rather than water. For example, using 1 gram of soil as the unit, the estimation of OTUs is considered to be between 2 000 and 5 000 (Schloss and Handelsman, 2005), and the number of distinct genomes, based on DNA reassociation kinetics, between 2 000 and 18 000 (Torsvik *et al*., 1990; Torsvik *et al*., 1996; Sandaa *et al*., 1999; Dunbar *et al*., 2001). Clearly, traditional microbiological methods for the isolation of microorganisms have the potential to determine only dominant populations, and tend to mask the detection of low-abundance species (Sogin *et al*., 2006). The large number of highly diverse, low-abundance species in an ecosystem constitutes a rare ‘biosphere’ that is largely unexplored. Recent developments in high throughput molecular methods are beginning to provide deep insights into a wide range of ecosystems, supplying information on microorganisms and their roles in the environment.

In conclusion, this study shows that the BCC of recycled irrigation freshwater is a diverse ecosystem, and supports the finding of Berkelmann *et al*. (1994), for which irrigation freshwater should be carefully preserved in order to avoid the colonisation of available ecological niches by single species and/or dangerous pathogens. In addition, in this study the use of a SSF showed that the BCC of irrigation freshwater can be altered by biofiltration, and that this had positive effect on plant growth. The use of fingerprinting methods such as T-RFLP provided useful information on the dynamics of bacterial populations in freshwater, and allowed rapid and inexpensive monitoring of the microflora. A consistant monitoring of recycled irrigation freshwater, however, remains of extremely importance, in order to avoid the intrusion of allochthonous microorganisms that can occur anytime in the distribution path of a hydroponic system (Hong and Moorman, 2005). Also, when dealing with closed hydroponic systems, this aspect becomes even more important: the invasion of external species has the potential of spreading to the entire crop, assuming that the environmental conditions are favourable for the pathogen.

## ACKNOWLEDGEMENTS

This study was part of the PhD work of Giovanni Cafà, funded by the University of Nottingham (Nottingham, UK) and the Food and Environment Research Agency (FERA; York, UK).

## REFERENCES

Alsanius, B.W., Nilsson, L., Jensen, P., Wohanka, W., 2001. Microbial communities in slow filters. Acta Hortic. 548, 591–601.

Altschul, S.F., Gish, W., Miller, W., Myers, E.W., Lipman, D.J., 1990. Basic local alignment search tool. J. Mol. Biol. 215, 403–410.

Anthony, R.M., Brown, T.J., French, G.L., 2000. Rapid diagnosis of bacteremia by universal amplification of 23S ribosomal DNA followed by hybridization to an oligonucleotide array. J. Clin. Microbiol. 38, 781–788.

Bashan, Y., Harrison, S.K., Whitmoyer, R.E., 1990. Enhanced growth of wheat and soybean plants inoculated with *Azospirillum brasilense* is not necessarily due to general enhancement of mineral uptake. Appl. Environ. Microbiol. 56, 769–775.

Baysal, O., Caliskan, M., Yesilova, O., 2008. An inhibitory effect of a new *Bacillus subtilis* strain (EU07) against *Fusarium oxysporum* f. sp. radicislycopersici. Physiological and Molecular Plant Pathology 73, 25–32.

Berkelmann, B., Wohanka, W., Krczal, G., 1995. Transmission of Pelargonium flower break virus (PFBV) by recirculating nutrient solution with and without slow sand filtration. Acta Hortic. 382, 256–262.

Berkelmann, B., Wohanka, W., Wolf, G.A., 1994. Characterization of the bacterial flora in circulating nutrient solutions of a hydroponic system with rockwool. Acta Hortic. 361, 372–381.

Brodie, E., Edwards, S., Clipson, N., 2002. Bacterial community dynamics across a floristic gradient in a temperate upland grassland ecosystem. Microb. Ecol. 44, 260–270.

Büttner, C., Marquardt, K., Führling, M., 1995. Studies on trasmission of plant viruses by recirculating nutrien solution such as ebb-flow. Hydroponics and Transplant Production 396, 265–272.

Calaway, W.T., Carroll, W.R., Long, S.K., 1952. Heterotrophic bacteria encountered inintermittent sand filtration of sewage. Sewage and Industrial Wastes 24, 642–653.

Calvo-Bado, L.A., Petch, G., Parsons, N.R., Morgan, J.A.W., Pettitt, T.R., Whipps, J.M., 2006. Microbial community responses associated with the development of oomycete plant pathogens on tomato roots in soilless growing system. J. Appl. Microbiol. 100, 1194–1207.

Calvo-Bado, L.A., Pettitt, T.R., Parsons, N., Petch, G.M., Morgan, J.A.W., Whipps, J.M., 2003. Spatial and temporal analysis of the microbial community in slow sand filters used for treating horticultural irrigation water. Appl. Environ. Microbiol. 69, 2116–2125.

Cardinale, B.J., 2011. Biodiversity improves water quality through niche partitioning. Nature 467, 86–89.

Chao, A., Bunge, J., 2002. Estimating the number of species in a stochastic abundance model. Biometrics 58, 531–539.

Chao, A., Chazdon, R.L., Colwell, R.K., Shen, T., 2006. Abundance-based similarity indices and their estimation when there are unseen species in samples. Biometrics 62, 361–371.

Clematis, F., Minuto, A., Gullino, M.L., Garibaldi, A., 2009. Suppressiveness to *Fusarium oxysporum* f. sp. radicis lycopersici in re-used perlite and perlite-peat substrates in soilless tomatoes. Biol. Control 48, 108–114.

Cole, J.R., Wang, Q., Cardenas, E., Fish, J., Chai, B., Farris, R.J., Kulam-Syed-Mohideen, A.S., McGarrell, D.M., Marsh, T., Garrity, G.M., Tiedje, J.M., 2009. The Ribosomal Database Project: improved alignments and new tools for rRNA analysis. Nucleic Acids Res. 37, D141–D145.

Colwell, R.K., 2006. EstimateS: statistical estimation of species richness and shared species from samples. Version 8. Department of Ecology & Evolutionary Biology, University of Connecticut, Storrs.

Cooper, A., 1979. The ABC of NFT. Grower books, London, UK.

Deniel, F., Renault, D., Tirilly, Y., Barbier, G., Rey, P., 2006. A dynamic biofilter to remove pathogens during tomato soilless culture. Agronomy for Sustanaible Development 26, 185–193.

Dunbar, J., Ticknor, L.O., Kuske, C.R., 2001. Phylogenetic specificity and reproducibility and new method for analysis of terminal restriction fragment profiles of 16S rRNA genes from bacterial communities. Appl. Environ. Microbiol. 67, 190–197.

Frenkel, O., Yermiyahu, U., Forbes, G.A., Fry, W.E., Shtienberg, D., 2010. Restriction of potato and tomato late blight development by sub-phytotoxic concentrations of boron. Plant Pathology 59, 626–633.

Garland, J.L., 1994. The structure and function of microbial communities in recirculating hydroponic systems. Advances in Space Research 11, 383–386.

Gertsson, U.E., Hansson, I., Waechter-Kristensen, B., Lundquist, S., Svedelius, G., Weich, R., 1994. Tomato grown in circulating nutrient solution using rockwool and as hydroponics. Acta Hortic. 361, 237–244.

Glick, B.R., 1995. The enhancement of plant growth by free-living bacteria. Can. J. Microbiol. 41, 109–117.

Hartmann, M., Widmer, F., 2008. Reliability for detecting composition and changes of microbial communities by T-RFLP genetic profiling. FEMS Microbiol. Ecol. 63, 249–260.

Hong, C., Moorman, G.W., 2005. Plant pathogens in irrigation water: challenges and opportunities. Crit. Rev. Plant Sci. 24, 189–208.

Horakova, K., Mlejnkova, H., Mlejnek, P., 2008. Evaluation of methods for isolation of DNA for polymerase chain reaction (PCR)-based identification of pathogenic bacteria from pure cultures and water samples. Water Science and Technology 58, 995–999.

Huisman, L., Wood, W.E., 1974. Slow sand filtration. World Health Organization, Geneva.

Huse, S.M., Huber, J.A., Morrison, H.G., Sogin, M.L., Welch, D.M., 2007. Accuracy of and quality of massively parallel DNA pyrosequencing. Genome Biology 8, R143.

Huson, D.H., Auch, A.F., Qi, J., Schuster, S.C., 2007. MEGAN analysis of metagenomic data. Genome Res. 17, 377–386.

Jakobsson, H.E., Jernberg, C., Andersson, A.F., Sjolund-Karlsson, M., Jansson, J.K., Engstrand, L., 2010. Short-term antibiotic treatment has differing long-term impacts on the human throat and gut microbiome. PLoS One 5.

Jensen, M.H., 1997. Hydroponics. HortScience 32, 1018–1021.

Joupert, E.D., Pillay, B., 2008. Visualization of the microbial colonization of a slow sand filter using an environmental scanning electron microscope. Electronic Journal of Biotechnology 11.

Kloepper, J.W., Lifshitz, R., Schroth, M.N., 1988. Pseudomonas inoculant to benefit plant production. ISI Atlas of Science: Animal and Plant Sciences, 30–34.

Koohakan, P., Ikeda, H., Jeanaksorn, T., Tojo, M., Kusakari, S., Okada, K., Sato, S., 2004. Evaluation of the indigenous microorganisms in soilless culture: occurence and quantitative characteristics in the different growing systems. Scientia Horticulturae 101, 179–188.

Lucy, M., Reed, E., Glick, R., 2004. Application of free living plant growth promoting rhizobacteria. Antonie Van Leeuwenhoek 86, 1–25.

McEniry, J., O'Kiely, P., Clipson, N.J.W., Forristal, P.D., Doyle, E.M., 2008. Bacterial community dynamics during the ensilage of wilted grass. J. Appl. Microbiol. 105, 359–371.

Menking, D.E., Emanuel, P.A., Valdes, J.J., Kracke, S.K., 1999. Rapid cleanup of bacterial DNA from field samples. Resources Conservation and Recycling 27, 179–186.

Muyzer, G., de Waal, E.C., Uitterlinden, A.G., 1993. Profiling of complex microbial populations by denaturing gradient gel electrophoresis analysis of polymerase chain reaction-amplified genes coding for 16S rRNA. Appl. Environ. Microbiol. 59, 695–700.

Muyzer, G., Teske, A., Wirsen, C.O., 1995. Phylogenetic relationships of Thiomicrospira species and their identification in deep-sea hydrothermal vent samples by denaturing grsadient gel electrophoresis of 16S rDNA fragments. Arch. Microbiol. 164, 165–172.

Nawrocki, E.P., Eddy, S.R., 2007. Query-dependent banding (QDB) for faster RNA similarity searches. PLOS Comput. Biol. 3, e56.

Omar, I., O’Neill, T.M., Rossall, S., 2006. Biological control of fusarium crown and root rot of tomato with antagonistic bacteria and integrated control when combined with the fungicide carbendazim. Plant Pathology 55, 92–99.

Page, D., Wakelin, S., van Leeuwen, J., Dillon, P., 2006. Review of biofiltration processes relevant to water reclamation via aquifers. CSIRO Land and Water.

Pagliaccia, D., Merhaut, D., Colao, M.C., Ruzzi, M., Saccardo, F., Stanghellini, M.E., 2008. Selective enhancement of the fluorescent pseudomonad population after amending the recirculating nutrient solution of hydroponically grown plants with a nitrogen stabilizer. Microb. Ecol. 56, 538–554.

Pares, R.D., Gunn, L.V., Cresswell, G.C., 1992. Tomato mosaic virus infection in a recirculating nutrient solution. J. Phytopathol. 135, 192–198.

Petri-Hansen, H., Steele, H., Grooters, M., Wingender, J., Flemming, H.C., 2006. Recent progress in slow sand and alternative biofiltration processes. IWA publishing, London, UK.

Pettitt, T.R., 2005. Slow sand filtration, a flexible, economic biofiltration method for cleaning irrigation water. A grower guide. Horticultural Development Council.

Postma, J., Lankwarden, J.B.L., van Elsas, J.D., 2001. Molecular fingerprinting of microbial populations in soilless culture systems. Acta Hortic. 548, 537–541.

Runia, W.T., van Os, E.A., Bollen, G.J., 1988. Disinfection of drainwater from soilless cultures by heat-treatment. Netherlands Journal of Agricultural Science 36, 231–238.

Sambrook, J., Fritsch, E., Maniatis, T., 1989. Molecular cloning: a laboratory manual. Cold Spring Harbor Press, New York.

Sandaa, R., Torsvik, V., Enger, Ø., Daae, F.L., Castberg, T., Hahn, D., 1999. Analysis of bacterial communities in heavy metal-contaminated soils at different levels of resolution. FEMS Microbiol. Ecol. 30, 237–251.

Schloss, P.D., Handelsman, J., 2005. Introducing DOTUR, a computer program for defining operational taxonomic units and estimating species richness. Appl. Environ. Microbiol. 71, 1501–1506.

Schloss, P.D., Westcott, S.L., Ryabin, T., Hall, J.R., Hartmann, M., Hollister, E.B., Lesniewski, R.A., Oakley, B.B., Parks, D.H., Robinson, C.J., Sahl, J.W., Stres, B., Thallinger, G.G., Van Horn, D.J., Weber, C.F., 2009. Introducing mothur: open-source, platform-independent, community-supported software for describing and comparing microbial communities. Appl. Environ. Microbiol. 75, 7537–7541.

Sogin, M.L., Morrison, H.G., Huber, J.A., Welch, D.M., Huse, S.M., Neal, P.R., Arrieta, J.M., Herndl, G.J., 2006. Microbial diversity in the deep sea and the underexplored "rare biosphere". Proc. Natl. Acad. Sci. U. S. A. 103, 12115–12120.

Stanghellini, M.E., Rasmussen, S.L., 1994. Hydroponics: a solution for zoosporic pathogens. Plant Dis. 78, 1129–1138.

Stecher, B., Chaffron, S., Käppeli, R., Hapfelmeier, S., Freedrich, S., Weber, T.C., Kirundi, J., Suar, M., McCoy, K.D., von Mering, C., Macpherson, A.J., Hardt, W., 2010. Like will to like: abundances of closely related species can predict susceptibility to intestinal colonization by pathogenic and commensal bacteria. PLOS Pathogens 6.

Torsvik, V., Goksoyr, J., Daae, F.L., 1990. High diversity in DNA of soil bacteria. Appl. Environ. Microbiol. 56, 782–787.

Torsvik, V., Sorheim, R., Goksoyr, J., 1996. Total bacterial diversity in soil and sediment communities - a review. J. Ind. Microbiol. 17, 170–178.

Waechter-Kristensen, B., Sundin, P., Berkelmann-Loehnertz, B., Wohanka, W., 1997. Management of microbial factors in the rhizosphere and nutrient solution of hydroponically grown tomato. International Symposium on Growing Media and Plant Nutrition 1, 335–340.

Wang, Q., Garrity, G.M., Tiedje, J.M., Cole, J.R., 2007. Naive Bayesian classifier for rapid assignment of rRNA sequences into the new bacterial taxonomy. Applied and Environmental Microbiology 73, 5261–5267.

Zhang, W., Tu, J.C., 2000. Effect of ultraviolet disinfection of hydroponic solutions on Pythium root and non-target bacteria. European Journal of Plant Pathology 106, 415–421.

